# Molecular Control of Circuit Plasticity and the Permanence of Imprinted Odor Memory

**DOI:** 10.1101/2022.03.29.486284

**Authors:** Yunming Wu, Limei Ma, Qiang Qiu, Wenjing Xu, Aviral Misra, Kyle Duyck, Jillian Blanck, Allison R. Scott, Shiyuan Chen, Huzaifa Hassan, Timothy J. Corbin, Andrea Moran, Kate Hall, Hua Li, Anoja Perera, C. Ron Yu

**Author notes:** Department of Biology, Howard Hughes Medical Institute, Stanford University, Stanford, CA 94305, USA.

## Abstract

Behavioral imprinting is a distinct form of learning that has a lifelong impact on social interactions and affectional behaviors^1-4^. Unlike other forms of memory, imprinting does not require conspicuous association of stimuli; exposure *per se* appears sufficient to induce memories that neither undergo extinction nor are altered by experience later in life. The site of storage of imprinted memory and the mechanisms that control its formation and permanence are unknown. Here we uncover a molecular mechanism that controls olfactory imprinting, which underlies behaviors including kin and nest recognition, maternal attachment, and homing^5-10^. We show that odor exposure during the perinatal period converts an innately aversive odor into a homing signal. The behavioral change is associated with odor-induced changes in the projection patterns of olfactory sensory neuron (OSN) expressing the cognate receptors for the exposed odor. We show that the Wnt signaling receptor Frizzled1 (Fzd1) acts as a master regulator of the critical period of OSN development and is responsible for closing the critical period to prevent further changes in the neural circuit. In Fzd1 knockout mice axon projection patterns are continually modified by sensory experience. As Fzd1 knockout abolishes the developmental critical period, it also abolishes odor imprinting. Specific knockout of Fzd1 in the OSNs have the same effect. Mechanistically, Fzd1 controls the critical period through an autoregulated shutdown and by controlling an activity-driven regulon in the OSNs. The transient expression and the subsequent downregulation of Fzd1 leads to the irreversible closure of the critical period to lock in circuits established during the critical period. The evidence suggests that imprinted odor memory is stored in the patterns of connectivity at the first synapse in the olfactory bulb. Early odor experience induces changes in the OSN projection to alter connectivity with innate circuits to establish a life-long memory.

## Introduction

In behavioral imprinting, brief experiences early in life have profound and life-long influences on animal’s behaviors. Young animals form bonds with parents, siblings, and their immediate birth environment based on sensory experience in early postnatal development^11^. Imprinting is distinct from other forms of memory in several aspects. Usually associated with attachment, it does not require overt pairing between the stimuli and reward signals. Pairing with negative stimuli, strikingly, can further enhance attachment rather than induce aversion^12, 13^. It is not clearly understood how imprinted memory is generated, where it is stored, what underlies the life-long memory that distinguishes it from other forms of learning, and how the acquisition is regulated at the cellular and molecular level. For imprinting to occur, the exposure must take place during a brief time window early in life. In studying the graylag geese following behaviors, Konrad Lorenz recognized that imprinting must occur at “a very specific physiological state in the young animal’s development”^1^. Finally, once formed, imprinted memories last a lifetime and do not appear to undergo extinction, nor are altered by experience later in life. These features raise possibility that behavioral imprinting is associated with the critical period in neural development^14, 15^. Moreover, the permanence of imprinted memory is reminiscent of genetically specified innate responses, raising the possibility that imprinting is to establish a connection between a stimulus with a hardwired behavioral circuit. To test these hypotheses, we establish a model of olfactory imprinting, identify the molecules involved in regulating the critical period of OSN development^16, 17^, and use this knowledge to manipulate the critical period, and examine the relationship between olfactory imprinting and odor-induced changes in olfactory circuit change.

### An olfactory imprinting paradigm

We first ask whether olfactory imprinting can be observed in mice. Odors are often described along an axis of pleasantness^18^. Postnatal odor exposure alters odor preference^12, 19-21^. To determine whether this change in preference fits the criteria of odor imprinting, we exposed neonatal pups to odors at various developmental periods and tested their behavioral responses at 3 months without further experience of the same odor (Fig. 1a)^22^. We focused on the use of acetophenone, an odor that triggered innate aversion (Fig. 1b). Mice exposed to acetophenone from birth till P7 exhibited preference rather than aversion as adults (Fig. 1b). The effect was odor-specific; exposure to acetophenone did not abolish aversion to another innately aversive odor, 2-phenylethylamine (PEA; Fig. 1c). Mice exposed for a shorter duration before P7, or after the P0-P7 window did not exhibit altered preference (Fig. 1b and d). Since the critical period of OSNs development is the first postnatal week^16, 17, 23^, these results were consistent with the notion that odor experience during the critical period resulted in odor imprinting.

**Fig. 1.**
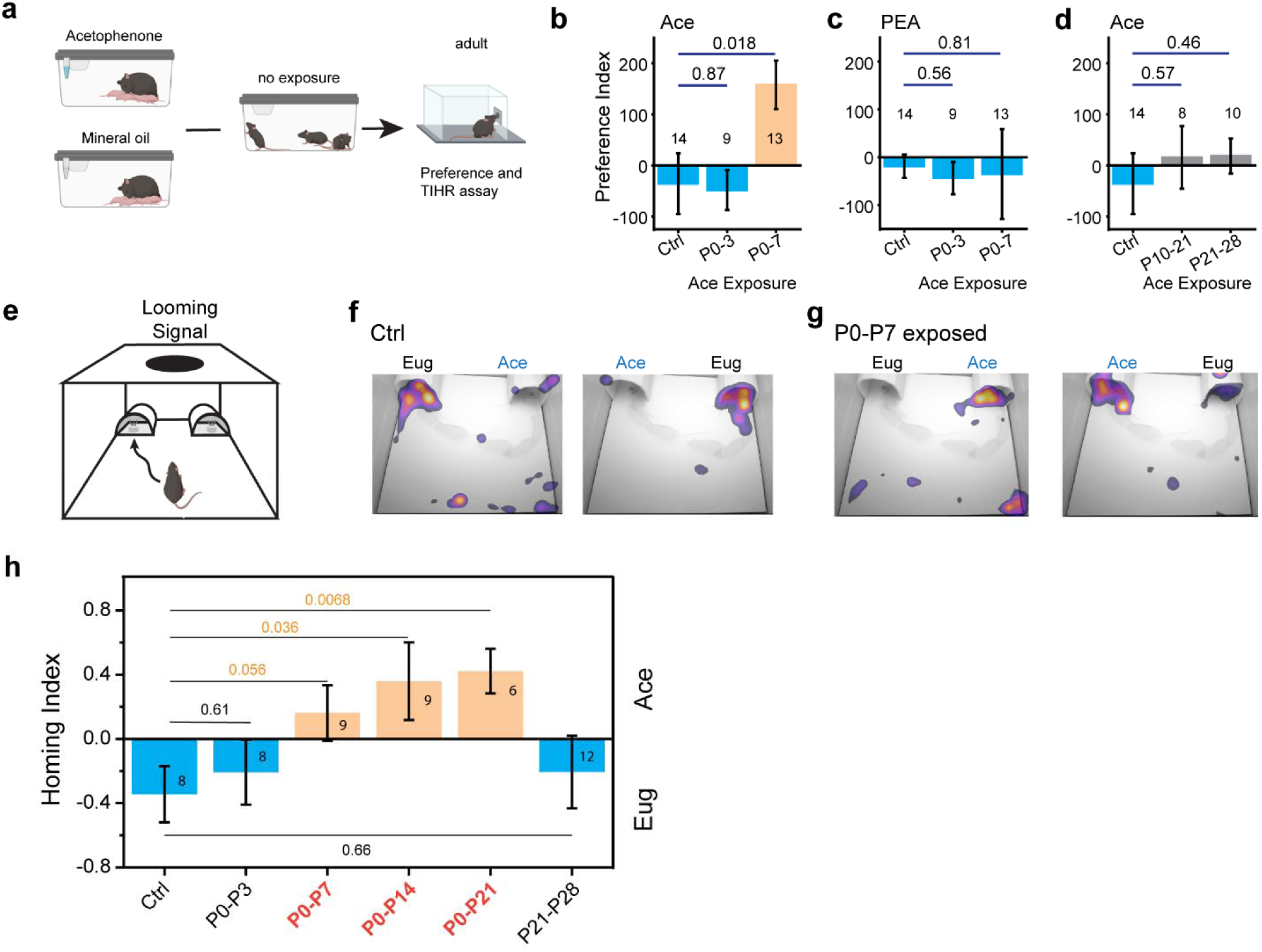
Olfactory imprinting turns an innately aversive odor into a homing signal. **a**. Schematic of experimental paradigm of assay for olfactory imprinting. Mice are exposed to an odor or mineral oil (carrier) during various postnatal periods (illustration shows neonatal exposure), followed by odor removal for an extended period and tested as adults for odor preference or threat induced homing response. **b**. Odor preference test of adult mice to acetophenone with no (control), P0-P3, or P0-P7 exposure to acetophenone. **c**. Same as (**b**) but tested for preference of 2-phenolethylamine (PEA), a predator odor that is innately aversive to mice. **d**. Same as (**b**) but acetophenone exposure took place during juvenile period. **e**. Illustration of the TIHR assay arena. **f**. Heatmaps showing control mice prefer igloos with the eugenol smell. **g**. Same as (**e**) but for mice exposed to acetophenone between P0-P7, which prefer igloos with the acetophenone smell. **h**. Quantitative measure of TIHR of different groups of mice. Mice exposed to acetophenone during the neonatal period strongly prefer acetophenone, but mice exposed during the P0-P3 or P21-P28 period do not. Numbers in the bars indicate the number of animals tested. One-way ANOVA was applied. *p* values are indicated above.

Odor preference assay can be complicated by familiarity, novelty seeking, and innate odor valence. Moreover, it did not provide an accurate measure for attachment associated with odor imprinting. A homing signal may not manifest as preference because it represents familiarity, comfort, and safety rather than pleasantness. We, therefore, further devised a threat induced homing response (TIHR) assay to evaluate attachment associated with imprinting (Fig. 1e; Extended data Fig. 1). We habituated animals in an open arena to allow free investigation of the arena and the two igloos placed at two corners. Following habituation, a looming signal ^24^ was triggered when the mouse wondered into a designated area at equal distance to the two igloos. We quantified the proportion of igloo in which the animal tried to hide. When the same odor was presented in both igloos, individual mice exhibited various tendency to hide under the igloos, but there was no overall preference (Extended Data Fig. 1). When a neutral (eugenol) and an innately aversive odor (acetophenone) were placed under separate igloos, mice strongly preferred the one with the neutral odor, even though there were no difference in the latency or duration of hiding (Fig. 1f; Extended Data Movie 1). In stark contrast, 3-months old mice that were exposed to acetophenone between P0-P7 exhibited a strong preference for the igloo with acetophenone over eugenol (Fig. 1g; Extended Data Movie 2). Even at 6 months of age, the animals had maintained preference towards acetophenone (Extended Data Fig. 3). Exposure beyond P7 induced stronger homing responses to igloos with acetophenone (Fig. 1h). Exposing the pups till P3, or from P21 to P28, did not change the homing preference. Thus, the TIHR assay revealed that odor experience during the critical period changed an innately aversive odor into a long-lasting homing signal.

### Precisely aligned timing between odor-elicited circuit and behavioral changes

Odor exposure also alters projection patterns of OSNs expressing the cognate receptors^22,25^. To determine whether these modifications provided a substrate for odor imprinting, we examined the projection patterns of OSNs expressing M71 (*Olfr151*) or M72 (*Olfr160*), both cognate receptors of acetophenone. We exposed the *Olfr151-IRES-tauGFP; Olfr160-IRES-tauGFP* mice (*M71G;M72G* mice for short) at birth, removed the odor at various time points, and evaluated the number of glomeruli innervated by M71 and M72 axons in adults (Figs. 2a-f)^26-28^. Odor exposure starting at birth led to divergent axon projection (Extended Data Fig. 4). Varying the starting point of odor exposure led to different numbers of GFP^+^ glomeruli observed at P21 (Fig. 2b-c). Exposure starting before P7 resulted in high numbers of glomeruli, with the peak at P3. There was a sharp drop in the number of glomeruli when odor exposure started after P7. Thus, there was a strong temporal correlation between odor-induced behavioral imprinting and changes in OSN axon projection patterns. The change was also odor-specific; exposure to acetophenone did not alter the projection patterns of axons expressing the MOR28 receptor (Extended Data Fig. 5).

**Fig. 2.**
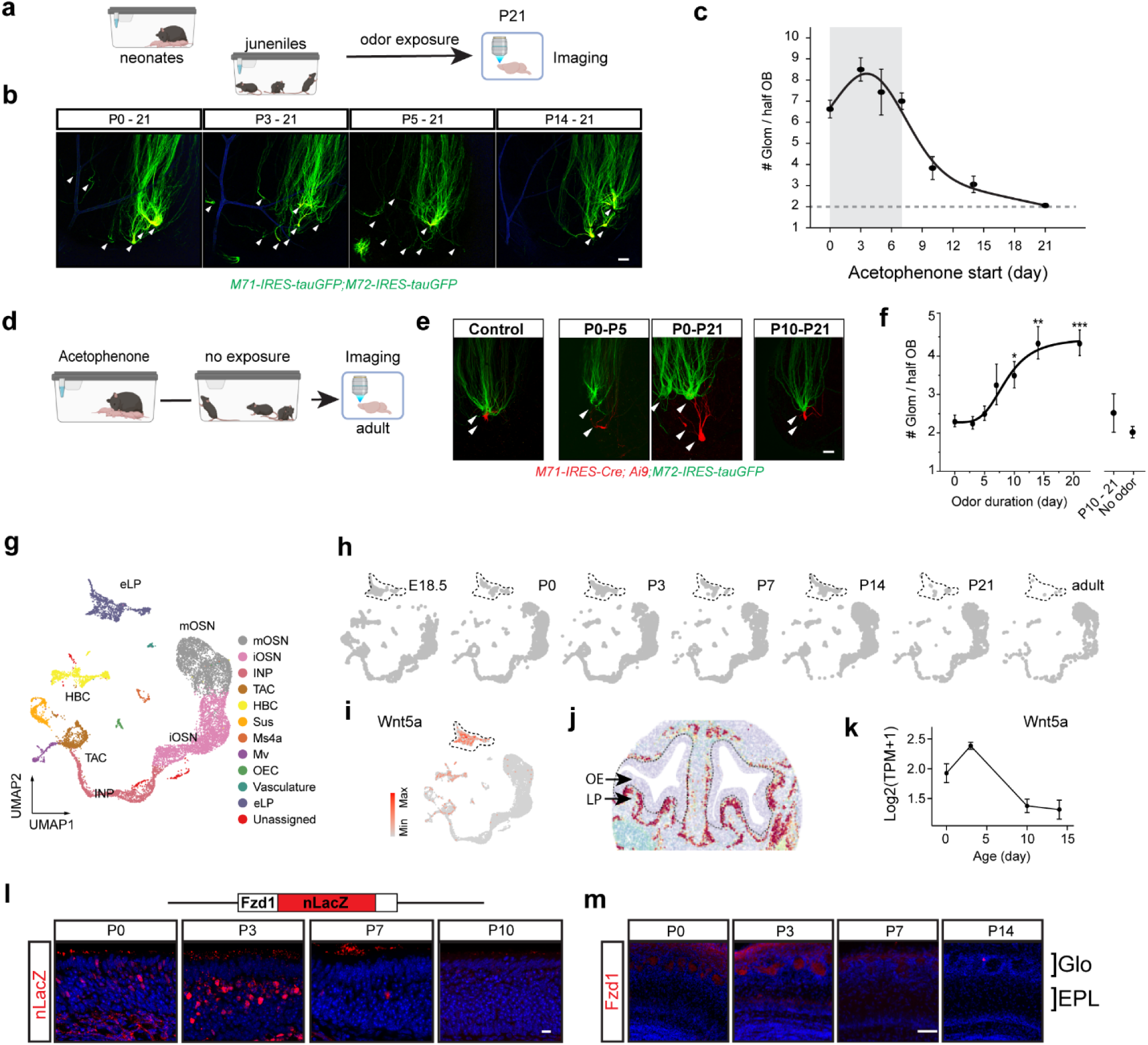
Coordinated expression of Wnt5a and Fzd1 coincides with odor induced changes in olfactory sensory axon projection. **a**. Schematic of experimental paradigm to examine the effect of odor exposure onset on axon projection. **b**. Sample images of axons labeled with GFP from *M71G; M72G* pups exposed to acetophenone at different start dates. Arrowheads indicate glomeruli containing the labeled axons evaluated from 3D confocal images. Scale bar: 100 µm. **c**. Quantification of the total number of M71 and M72 glomeruli at P21. Shaded area indicates the period during which odor exposure causes supernumerary projections. **d**. Schematic of experimental paradigm to examine recovery of supernumerary axon projection following odor removal. **e**. Sample images of axons labeled with tdTomato and GFP from *M71R; M72G* pups. **f**. Quantification of the total number of M71 and M72 glomeruli at adulthood. Shaded area indicates the period when axon can remodel to restore single glomerular projection. **g**. UMAP visualization of cells from the scRNA-Seq experiment. Color code indicates cell type. INP, intermediate neuronal progenitor. TAC, transient amplifier cell. HBC, horizontal basal cell. Sus, sustentacular cell. Ms4a, cells expressing Ms4a receptors. Mv, Microvillous cell. OEC, olfactory ensheathing cell. eLP, early lamina propria cells. **h**. The same UMAP in (g) plotted by age. Dashed line circles the eLP cells. **i**. Expression of Wnt5a determined by scRNA-Seq. Data is presented as SC transformed values. Dashed line circles the eLP cells. **j**. Locations of eLP cells as predicted by Slide-Seq in a P3 OE section. Dashed line demarks the boundary between the olfactory epithelium (OE) and lamina propria (LP). **k**. Expression profile of Wnt5a during postnatal development determined by bulk RNA-Seq. Data is presented as log2 (TPM + 1). **l**. Expression of Fzd1 analyzed in *Fzd1*^*+/nlacZ*^ mouse line using antibody against lacZ (red). Scale bar, 10 µm. **m**. Fzd1 protein the olfactory bulb stained with anti-Fzd1 antibodies (red). Glomerular (Glo) and external plexiform layer (EPL) of the olfactory bulb are marked. Sections counter stained with DAPI (blue). Scale bar, 50 µm.

We next exposed the neonatal pups to acetophenone and removed the odor at various timepoints and examined the glomeruli innervation in adult (Fig. 2d). Upon odor removal, the number of *M71G;M72G* axons was comparable with no odor controls if odor exposure was limited to P7 or earlier (Figs. 2e and f). This observation suggested that the axons were able to reconverge following odor removal. However, exposure beyond P7 resulted in supra-numeric innervation (Figs. 2e and f). This result indicated that the divergent projection patterns resulted from odor exposure were imprinted after the end of the critical period^16, 17^. Taken together, odorant exposure triggered axon rewiring during the first week, after which the projection pattern was secured following the closure of the critical period.

### Coordinated expression of Wnt5a and Fzd1 during the critical period

The association between OSN axon re-wiring and olfactory imprinting suggested that the anatomical changes may be responsible for the behavioral response. To further investigate this association, we sought to identify the molecules that control the critical period. We reasoned that a signaling mechanism might heightened the plasticity of OSN axons and sought differentially expressed genes specifically during the critical period. In single cell RNA-Seq (scRNA-Seq) of olfactory epithelia (OE) from embryonic day 18.5 (E18.5) to adult (Fig. 2g), we have identified most known cell types in the OE whose distribution was relatively stable across the ages. In contrast, a prominent population of cells during early development disappeared after P14 (Fig. 2h). These cells expressed Wnt5a (Fig. 2i) and were located in the lamina propria using Slide-Seq^29^, just underneath the sensory epithelia where OSN axons exit (Figs. 2j; Extended Data Fig. 6). These cells expressed connective tissue markers including several collagen genes, but also genes that were unusual for connective tissues, including the non-coding RNA H19 and several Igf binding proteins. They were distinct from the olfactory ensheathing cells, which were mostly located near the olfactory bulb (Extended Data Fig.6). These, which we named the early Lamina Propria (eLP) cells, were likely chondrocytes responsible for cartilage formation during early period. Consistent with scRNA-Seq, bulk RNA-Seq detected Wnt5a expression during first postnatal week, with the highest level at P3 (Fig. 2k). We also identified several Wnt receptors expressed in the OE, but only Fzd1 exhibited heightened expression in the first postnatal week. Using a mouse line in which Fzd1 coding region was replaced by nuclear lacZ from the Fzd1 allele (*Fzd1*^*+/nlacZ*^)^30^, we confirmed that Fzd1 was expressed by the OSNs, with the highest level at P3 (Fig. 2l)^31^. Fzd1 protein was also prominently detected in the olfactory glomeruli at P3 and P7, but not later (Fig. 2m). Thus, Wnt5a and Fzd1 are coordinately expressed during the critical period, raising the possibility that their interaction enables plastic changes in axon projection in that time window.

### Fzd1 is required for the closure of the critical period

Wnt5a has been shown to serve as a morphogen to promote axon growth^32^. We examined OSN axon projections in homozygotic *Fzd1*^*nlacZ/nlacZ*^ mice^30^. Surprisingly, besides the observation of occasional stray axons, the projection patterns the OSNs were indistinguishable between *Fzd1*^*nlacZ/nlacZ*^ mice and control (Figs. 3a and 3b). Thus, Fzd1 did not serve as a receptor of morphogens for OSN targeting as proposed for CNS neurons. On the other hand, we observed a striking difference in GFP^+^ glomeruli when *M71G; M72G*; *Fzd1*^*nlacZ/nlacZ*^ pups were exposed to acetophenone at different postnatal dates (Fig. 3c; Extended Data Fig 7). Odor exposure during the first week also led to projection into ectopic glomeruli, but the number of GFP^+^ glomeruli in *Fzd1*^*nlacZ/nlacZ*^ mice that began exposure at P3 was significantly lower than in controls (Fig. 3c). Strikingly, in *Fzd1*^*nlacZ/nlacZ*^ mutant, acetophenone exposure at P10 and P14 continued to result in supernumerary glomeruli innervation, indicating that odor exposure past the critical period still drove axon divergence.

**Fig. 3.**
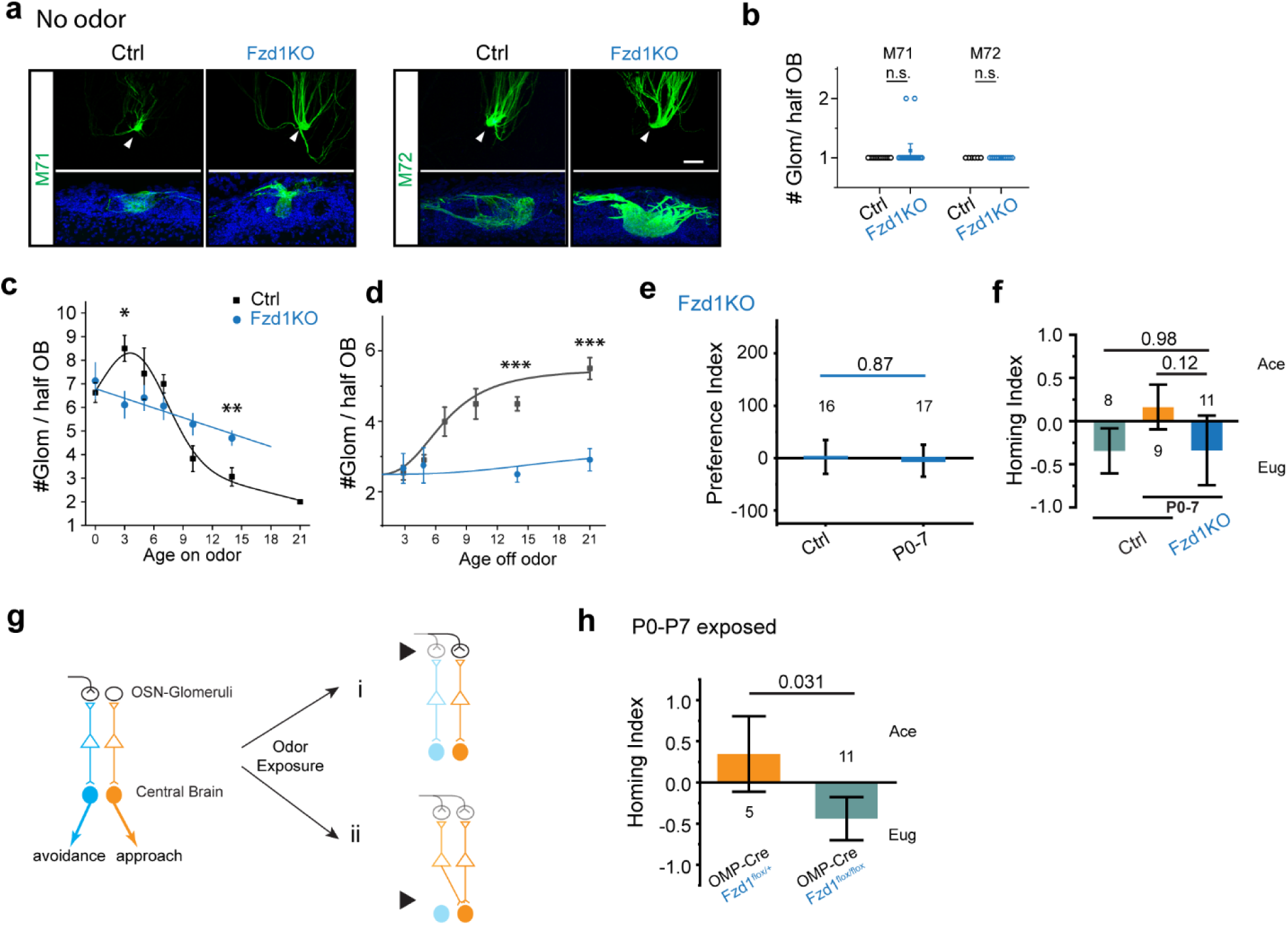
Fzd1 is required for closure of critical period of OSN development and imprinted odor memory. **a**. Projection patterns of M71 and M72 axon in the control and *Fzd1*^*nlacZ/nlacZ*^ (*Fzd1KO*) mice without odor exposure. Upper panels, representative images of whole mount images of the dorsal OB. Lower panels, OB sections stained with antibody against GFP (green) and DAPI (blue). **b**. Quantification of the M71and M72 glomeruli from control (black, n = 17 and 8 for M71 and M72 glomeruli, respectively) and *Fzd1KO* (blue, n = 19 and 11) animals. Circles represent individual data points. **c** and **d**. Quantification of the effect of acetophenone exposure onset (c) or offset (d) on axon projection in control and *Fzd1KO* mice. Experiments were conducted as figure 2a-f. Squares (control) and filled circles (*Fzd1KO*) represent mean. Control curves are the same as figure 2c and 2e, respectively. **e**. Effect of neonatal acetophenone exposure on adult odor preference of *Fzd1KO* mice **f**. Result of TIHR assay for *Fzd1KO* mice, which no longer exhibit homing towards igloos with acetophenone. **g**. Diagram of two general hypotheses. Left: two circuits mediating innate odor preference (orange) or aversion (blue). Right: two scenarios imprinting may change behaviors. Top (i): neonatal odor exposure leads to wiring changes such that OSNs recognizing the odor are connected to the approach pathway. Bottom (ii): neonatal odor exposure leads to changes in central brain to create association between odor and approach behaviors. The two scenarios are not mutually exclusive. **h**. Testing the first hypothesis by OSN-specific Fzd1 knockout. Homing indices for control and knockout mice indicate that OSN-specific knockout is sufficient to abolish changing an aversive odor into a homing signal. Significance test was performed with one-way ANOVA. Number of animals and *p* values are indicated. n.s., not significant (*p* > 0.05)

In the converse experiment where the pups were exposed to acetophenone starting from P0 with the odor removed at different postnatal days to allow axons to recover, the M71 and M72 axons in *Fzd1*^*nlacZ/nlacZ*^ mice converged back to two glomeruli per half bulb even when we delayed odor removal till P14 and P21, compared with an average of ∼6 glomeruli in the controls (Figs. 3d; Extended Data Fig. 8). This result indicated that in the *Fzd1*^*nlacZ/nlacZ*^ mice remodeling of axon projections continued beyond the normal critical period. We further confirmed the extended plasticity using genetic perturbation of OSN projections using the OMP-tTA; tetO-Kir2.1-IRES-tauLacZ mice (Extended Data Fig. 9)^17, 33^. Thus, the *Fzd1*^*nlacZ/nlacZ*^ mice exhibited continuous plasticity of axon remodeling. The expression Fzd1 was required to terminate the plasticity and close the critical period. In the absence of Fzd1, the axon projection can be continuously altered by the environment or restore the default projection pattern in the absence of continuous odor stimulation.

### Fzd1 is required for imprinted olfactory memory

That Fzd1 knockout abolished the critical period of OSN development provided an opportunity to further test whether odor imprinting required the critical period. We thus examined odor preference of *Fzd1*^*nlacZ/nlacZ*^ mice, with or without early postnatal exposure (Fig. 3e and f). As adults, the *Fzd1*^*nlacZ/nlacZ*^ exhibited the same level of aversion regardless of neonatal exposure took place (Fig. 3e). In the TIHR assay, we also found that *Fzd1*^*nlacZ/nlacZ*^ mice exhibited no preference to acetophenone (Fig. 3f). These results indicated Fzd1 was required for odor imprinting. A plausible explanation was that without the closure of the critical period, the continuous remodeling of axon projection removed the memory trace that would otherwise have been preserved by securing the altered connection.

These experiments established a strong connection between odor experience induced change in axon projection and odor imprinting. One interpretation of the results is that imprinted memory is stored in the patterns of OSN to glomeruli projection (Fig. 3g, i), where the glomerular set representing the odor is wired to a circuit mediating the approach behaviors. Alternatively, odor exposure may induce changes along the olfactory pathway in the brain (Fig. 3g, ii). Fzd1 expression in other parts of the brain and may control a critical period not associated with the OSNs. To test whether the critical period in OSN development *per se* is associated with imprinted memory, we engineered a conditional deletion allele of Fzd1 by flanking the Fzd1 coding region with two loxP sites. Crossing the Fzd1^flox/flox^ line with the OMP-IRES-Cre line allowed us to specifically knockout Fzd1 in the OSNs. Exposure to acetophenone between P0-P7 elicited a preference for acetophenone in the TIHR assay in the control OMP-IRES-Cre;Fzd1^flox/+^, but not OMP-IRES-Cre;Fzd1^flox/flox^ mice (Fig. 3h). These results indicated a cell-autonomous effect of Fzd1 in regulating the critical period of the OSNs and the plasticity of experience-induced axon projection *per se* is associated with imprinted memory.

### Auto-regulation by Fzd1 leads to closure of critical period

Fzd1-mediated Wnt signaling can promote axon growth, but it seemed paradoxical that the transient expression of Fzd1 would lead to the closure of the critical period to prevent further circuit modification. We hypothesized the Fzd1 expression formed a self-inhibitory loop to autoregulate itself. To test this hypothesis, we generated a mouse line that ectopically expressed Fzd1 under the tetracycline promoter (*tetO-Fzd1-IRES-tTomato*; Extended Data Fig. 10). In compound heterozygotic *OMP-IRES-tTA;tetO-Fdz1-IRES-tdTomato; Fzd1*^*+/nlacZ*^ (*Fzd*^*EE*^*;Fzd1*^*+/nlacZ*^) mice, exogenous Fzd1 was induced to be expressed in the mature OSNs. In the meantime, we could monitor the expression from the endogenous *Fzd1* locus by examining nuclear LacZ (Fig. 4a and b). In *Fzd1*^*EE*^ mice, endogenous Fzd1 expression was terminated by P1, indicating that it was inhibited by ectopic Fzd1 expression (Fig. 4b). DOX administration turned off expression from the transgenic allele (Extended Data Fig. 11) but did not restore expression from the endogenous locus (Fig. 4c). Thus, Fzd1 permanently downregulated its own expression.

**Fig. 4.**
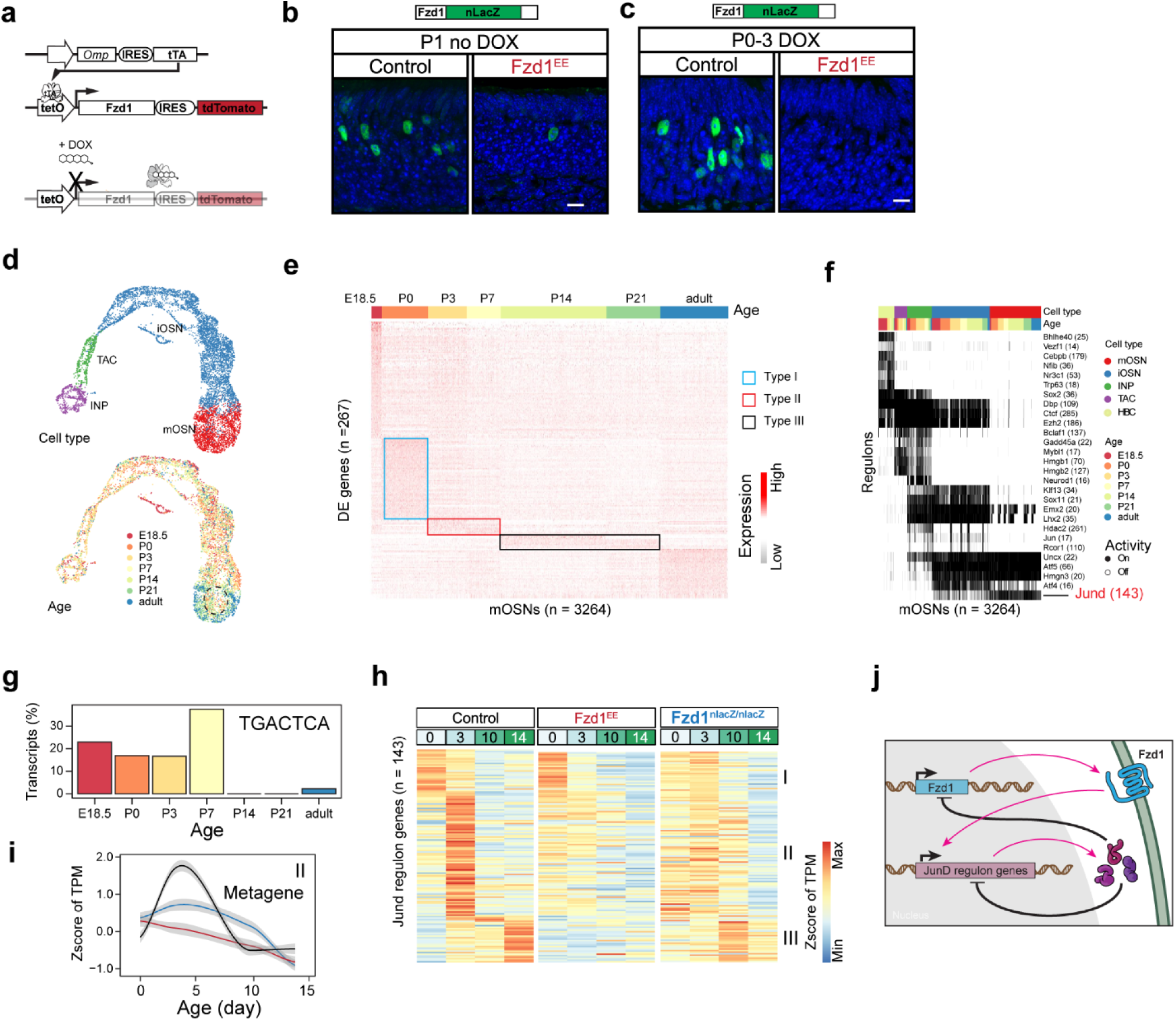
Autoregulation of Fzd1 controls dynamic gene expression in the critical period. **a**. Illustration of strategy to induce ectopic expression of Fzd1 in the mOSNs using the tet-off system. **b**. Ectopic Fzd1 expression in OMP-IRES-tTA;tetO-Fzd1-IRES-tdTomata (*Fzd1*^*EE*^) mince suppresses endogenous Fzd1 as early as P1. Endogenous Fzd1 was assessed as nLacZ expression. **c**. Dox administration suppressed ectopic Fzd1 expression but did not restore endogenous Fzd1 expression. **d**. UMAP representations of single cells of the OSN lineage and color coded according to cell type (top) or age (bottom). Dashed line circle mOSNs in early stages. **e**. Heatmap showing genes differentially expressed during development in mOSNs. Boxes indicate three groups of genes expressed between P0 and P21. Data is represented as SCT. **f**. Regulon analysis using genes differentially expressed during development. Each column is one cell. Each row shows the presence of one regulon in individual cells. Regulon activity was binarized. JunD regulon was highlighted in red. **g**. The percentage of genes in JunD regulon containing “TGACTCA” motif within 500 bp window flanking the transcription start site grouped by the time of peak expression. **h**. Heatmap showing the expression profile of genes in JunD regulon during development in control, *Fzd1*^*EE*^, and *Fzd1*^*nlacZ/nlacZ*^ OEs. Data is presented as Z-score of the Transcripts Per kilobase Million (TPM). **i**. Metagene analysis of the Type II genes in the JunD regulon. Metagene is calculated as a locally estimated scatterplot smoothing (LOESS) of genes within the group. Shaded area indicates 0.95 confidence interval. **j**. Schematic illustration of a model of Fzd1 action. Fzd1 triggers the JunD regulon expression, which in turn have negative feedback to turn off Fzd1 and the regulon genes.

We next determined whether Fzd1 exerts a broader transcriptional control in the OSNs. Bulk RNA-Seq showed a sharp transition of gene expression profile in the OE at P7 when the Fzd1 expression ended, and the critical period was closed (Extended Data Fig. 12). To identify genes that controls the critical period in the OSNs, we further analyzed the differentially expressed genes in the scRNA-Seq data in the OSN lineage (Fig. 4 d-f). We plotted the UMAP representation according to age of the animal and found that the mature OSNs (mOSNs) segregated according to age (Fig. 4d) and subsets of genes were specifically associated with different developmental stages (Fig. 4e). Using SCENIC^34^,we identified several regulons controlled by known transcription factors associated with specific cell types in the OE (Fig. 4f). Four regulons were found in the mOSNs, but only the JunD regulon was specific to mOSNs alone (Fig. 4f). It contained 143 genes, many of which were regulated by neuronal activity or by calcium dependent pathways, including Jun, JunD, Cebpb, and the cAMP responsive transcription factor ATF4. Calcineurin, calmodulins, calmodulin dependent kinases, immediate early genes, as well as several axon guidance molecule genes were part of this regulon. A significant number of genes expressed at early time points (E18.5-P7) contained the “TGACTCA” motif within 500 bp region flanking the transcription start sites (Fig. 4g). This motif is a known binding site for activity dependent AP-1 transcription factors c-Fos and c-Jun. JunD itself is a member of AP-1 transcription factor, which can assemble into newly synthesized NFAT transcription complex upon activation^35^. In neurons, NFAT signaling can be activated by receptor tyrosine kinase or GPCR and have been associated with axon remodeling of OSNs in zebrafish and auditory critical period^36-41^. This regulon could be the target of Fzd1 activation for transcriptional control. Although the canonical Wnt pathway genes are known to regulate gene expression, they are not detected in the OSNs as indicated by our bulk and single cell RNASeq experiments.

We next tested whether Fzd1 regulated the JunD regulon during the critical period through RNASeq analysis of differentially expressed genes in knockout (*Fzd1*^*nlacZ/nlacZ*^) and ectopic expression (*Fzd1*^*EE*^) mice (Figs. 4h-i). During postnatal development, there were three broad temporal profiles in gene expression (Extended Data Fig 12). Type I and III genes exhibited continuous decline (I) or increase (III) after birth. Type II genes were detected at P0, peaked at P3, and decreased after P7. GO term analysis indicated that the Type II group was enriched in genes controlling cell fate and chromosome organization (Extended Data Fig 12). The three types were also observed for the genes within the JunD regulon (Fig. 4b and e). The Type II genes had a dynamic expression profile associated with the critical period. Their expressions were significantly suppressed in *Fzd1*^*EE*^ but elevated in the *Fzd1*^*nlacZ/nlacZ*^ OE (Figs. 4h-i). Thus, Fzd1 negatively regulated of the expression of these genes. Moreover, these genes did not have a peak expression at P3 in the *Fzd1*^*nlacZ/nlacZ*^ OE, suggesting that the induction of their expression also required Fzd1 (Fig. 4i). Several nucleotide interacting factors involved in cell fate determination, including Uncx, Msi2, Jarid2, Ebf1, and Ebf3 in the JunD regulon responded to Fzd1. They provide a potential link between Fzd1 activation through G-proteins and the downstream transcriptional controls. Taken together, the evidence suggests a model that the transient expression of Fzd1 provides a feedback signal to downregulate Fzd1 itself and the regulon genes (Fig. 4j). This downregulation could lead to the termination of the critical period. In Fzd1 knockout, the induction and suppression do not occur, and the plasticity persists.

## Discussion

At birth, animals are endowed with sensory circuitries not only to sense the environment, but also elicit innate responses. Neural circuits mediating innate responses to sensory stimuli are thought to be genetically hardwired. Increasing evidence show that sensory experience can alter the processing of these sensory cues. Here we show that early odor experiences can turn an innately aversive odor into a homing signal. Moreover, odor experience during the critical period specifically changes the projection pattern of OSNs expressing the cognate receptors. Our study revealed a pivotal role of Fzd1 in regulating the critical period, and that Fzd1 signaling mediated by the JunD regulon triggers a cascade of transcriptional events that lead to the shutdown of the critical period. Genetically abolishing the critical period by knockout Fzd1 prevented olfactory imprinting. Since imprinted odor memory shares similarity with innate odor responses in that the responses are impervious to extinction or other sensory experiences, these results suggest a model that olfactory imprinting alter the valence associated with the odors through experience-dependent rewiring of sensory input. This model is akin to invertebrate brains that possess labeled lines to mediate approach/avoidance behaviors. In *C. elegans*, for example, expression of an odorant receptor in its normal set of olfactory neurons allows the animal to chemotaxis, but its misexpression in different neurons leads to avoidance^42^. The site (neurons) of receptor expression determines the valence of the stimulus. Our study suggests that changing the wiring diagram at the early processing stage can lead to switches in the behavioral responses. By tying changes in the wiring diagram to the critical period, it permits a window of flexibility for the behaviors to adapt to the natural environment while preserve the permanence of the innate circuit. The closure of the critical period prohibits further modification of the connection to secure established association between the input patterns and the behavioral circuits. An alternative scenario that the preference circuit develops earlier than the avoidance circuit, and early experience permit the association between an odor and the approach behavior. However, this model does not explain the strict requirement of the critical period in OSN development for odor imprinting.

## Acknowledgments

We thank Dr. Ai Fang, Troy Green, Andrew Box, Lab Animal Service Facility, Microscopy Center, and Tissue Culture Core of Stowers Institute for their technical assistance, Dr Fei Chen and Mr. Evan Murray for sharing reagents and experiment protocols of SLIDE-Seq2. We also thank Yu lab members for their insightful discussions. This work was supported by funding from NIH (R01DC016696 and R01DC014701) and Stowers Institute for Medical Research to C.R.Y.

## Author Contributions

Conceptualization, C.R.Y. and Y.W.; Investigation, Y.W., L.M., Q.Q., A. Misra. J.B., Y.Z., A.R.S., W.X.; Methodology, Y.W., A.V.; Transgenic Mice: Y.W., L.M., A. Moran., T.J.C. Software, S.C., A. Misra, Q.Q.; Formal Analysis: Y.W., Y.Z., Q.Q, and H.H.; Validation, K.D.; Writing – Original Draft, C.R.Y. and Y.W.; Writing – Review & Editing, C.R.Y., L.M., and Y.W.; Supervision, C.R.Y., A.P., H.L.; Funding Acquisition, C.R.Y.

## RESOURCE AVAILABILITY

### Lead Contact

Further information and requests for resources and reagents should be directed to and will be fulfilled by the Lead Contact, C. Ron Yu.

### Materials Availability

The plasmids used in this study are available from Addgene. Transgenic mice generated in this study will be available from the Jackson Laboratory.

### Data and code availability

Proteomic and RNA-Seq data in this study are submitted to the public data repositories detailed in the key resource table. Other original data underlying this manuscript can be accessed from the Stowers Original Data Repository at: http://www.stowers.org/research/publications/libpb-1501

## Material and Methods

### Experiment animals

Animals used in this study are described in the key resource table. All animals were maintained in Lab Animal Services Facility of Stowers Institute with a 14:10 light cycle and provided with food and water *ad libitum*. Experimental protocols were approved by the Institutional Animal Care and Use Committee at Stowers Institute and in compliance with the NIH Guide for Care and Use of Animals. The genotypes of the animals were determined by Transnetyx and in-house PCR.

### Transgenic mice

For the tetO-Fzd1-IRES-tdTomato construct, the Fzd1 coding region was cloned from plasmid pRK5-mFzd1, a gift from Chris Garcia & Jeremy Nathans (Addgene #42253), into MluI and PacI sites of the plasmid tetO-V1rj3-IRES-tdTomato ^43^ using Gibson assembly. The fragment between AscI and FseI sites was digested and purified through size exclusion chromatography using Toyopearl HW-75 resin (Tosoh Bioscience). The purified fragments were injected into through standard pronuclei injection at the LASF of Stowers Institute.

### Odor stimulation

Pups were fostered to a CD-1 mom before the odorant stimulation. 1.5 mL of acetophenone (Millipore Sigma) was placed a microcentrifuge tube over the cage top beyond the animals’ reach. The odorant was refilled daily to maintain the same volume over time.

### Odor preference assay using PROBES

Hardware design files, parts list, software source code, compiled program and instruction manuals of PROBES are published previously ^44^ and can be found at the Stowers Institute FTP site, ftp://ftp.stowers.org/pub/yu_lab/PROBES/. Acetophenone (Millipore Sigma) was diluted into mineral oil at 1:10^3^ (v/v). Animals were tested with 4 trials of mineral oil and 4 trials of odor, with 5 minutes in each trial. Preference score was defined as:

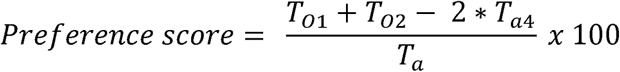

where *T*_*O*l_ is the sum of poke duration in the first odor trial. *T*_*O*2_ is the sum of poke duration in the second odor trial. *T*_*a*4_ is the sum of poke duration in the fourth air trial. *T*_*a*_ is the average of the 4 summations of poke duration in the air trials.

### Threat induced homing response assay

The TIHR assay was modified from looming induced escape response assay^24^. The test arena was a 40cm (long) × 40cm (wide) × 30cm (height) box with plexiglass walls and bottom. Half-cylindrical ‘igloos’ with the dimension of 10cm (long) × 10cm (wide) × 7.5cm (height) were manufactured using 3-D printing. Two igloos were placed along one wall on each corner, with the openings facing away from the wall. A computer monitor with pure white background was placed on the top of the arena. It covered approximately 70% of the space above the arena, leaving a gap between the monitor and one edge of the box to allow video recording. Mouse movement was recorded using a wide-angle USB camera at 1920 × 1080 resolution and 30fps. In a typical experiment, a mouse was placed in the arena to habituate for 30 minutes. After habituation, the arena was cleaned, and two new igloos were placed at the two corners. Cotton bedding pieces (1cm × 1cm) soaked with 1 ml odors (acetophenone or eugenol at 1:1000 dilution in mineral oil) were put at the centers of entrance of the igloos. The mouse was allowed to investigate the igloos for 2 minutes before the looming stimulus program was activated. Looming signal generation and delivery followed a published study^45^. The signal was an expanding dark disc that eventually cover the entire screen. A closed-loop workflow was written in Bonsai^46^ with the BonVision package^47^ to control the stimulation. As the mouse entered a specified region of interest (ROI) that was equidistant to both the igloos, the looming stimulation was triggered. Each stimulation epoch consisted of 20 repeats of consecutive looming sweeps, with each sweep consisted of an expanding signal that took 250ms to cover the entire screen, followed by 500ms darkness. A 10 second timeout was imposed before another stimulus could be given. The entire session last 15 minutes.

Mouse posture and movement were tracked using the DeepLabCut software^48^. The nose position was used to calculate the dwell time at different parts of the arena. Custom MATLAB codes were written for image correction for lens distortion, image registration, and calculating movement trajectory, heatmap, latency (time for mouse to enter the igloo region) and duration (time spent in igloo region). Homing index was calculated as:

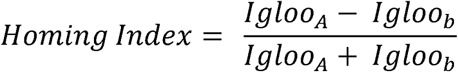

*Igloo*_*A*_ and *Igloo*_*B*_ are the number of times the animal escape into the A and B igloo, respectively.

### Spectrum imaging of the axon projection

The imaging of the glomeruli was conducted using a spectrum imaging method described previously ^23^. Animals were sacrificed by cervical dislocation. The skin and skull above the dorsal glomeruli were removed. For the MOR28 glomeruli, the olfactory bulb was dissected. The tissue was washed in 10 mM PBS 3 times to remove blood. After washing, the sample was mounted onto an imaging dish with No. 1.5 coverglass (Cellvis) and imaged using LSM780 confocal microscope (Zeiss) equipped with QUASAR detector. The spectral separation was conducted in Zen using linear unmixing.

### Immunohistochemistry

The olfactory bulb (OB) immunohistochemistry was conducted using free-floating sections. The animals were anesthetized with urethane (2 mg / g body weight). Tissue was fixed by intracardial perfusion with 10 mL 10 mM PBS, followed with 10 mL 4% paraformaldehyde (PFA) in PBS. The olfactory bulb was dissected and post-fixed with 4% PFA in PBS at 4°C overnight. After the fixation, the samples were sectioned using a vibratome (Leica VT1000), stained with primary antibodies in PBSTD (10 mM PBS, 0.1% Triton X-100, 3% donkey serum) overnight at room temperature with gentle agitation. Then the sections were washed with PBST (10 mM PBS, 0.1% Triton X-100) 5 minutes for 3 times, stained with secondary antibodies and DAPI in PBSTD for overnight at room temperature with gentle agitation. After washed with PBST 5 minutes for 3 times, the sections were mounted onto slides with No. 1.5 coverslip using Y-mount for imaging^23^.

The olfactory epithelium (OE) immunohistochemistry was conducted using cryo-sections. After post-fixation, the samples were de-calcified in 0.5 M EDTA with 30% sucrose at 4°C overnight, embedded in O.C.T., snap frozen in liquid nitrogen, and stored in -70°C until use. The tissue blocks were cut into 10 µm sections using a cryostat (CryoStar NX70) and mounted on charged slides. The sections were dried on a slide warmer at 100□ for 2 minutes, stained with primary antibodies in PBSTD overnight at room temperature with gentle agitation. Then the sections were washed with PBST 5 minutes for 3 times, stained with secondary antibodies and DAPI in PBSTD for overnight at room temperature with gentle agitation. After washed with PBST 5 minutes for 3 times, the sections were mounted with No. 1.5 coverslip using Y-mount for imaging.

Both OB and OE immunohistochemistry images were taken using LSM700 confocal microscope (Zeiss). Antibodies used in this study were listed in key resource table. Image cropping and contrast enhancement were conducted in Fiji ^49^.

### Bulk RNA-Seq of the olfactory epithelium

For the RNA-Seq experiment described in Figure 2 and S2, the OE were dissected from the nasal cavity directly into Trizol (Thermo Fisher Scientific). RNA was extracted according to manufacturer’s instruction. Sequencing libraries were generated using the mRNA-Seq Sample Prep Kit (Illumina) and sequenced individually as 75 bp single end reads on Illumina Genome Analyzer II system at a depth of 123-161 thousand clusters per sample.

For the time course RNA-Seq of *Fzd1*^*nlacZ/nlacZ*^, *Fzd1*^*EE*^, and their littermate control animals, the OE were dissected directly into Trizol. RNA was extracted using Direct-zol RNA purification kit (Zymo Research) according to manufacturer’s instruction. Libraries were made using TruSeq Stranded Total RNA Library Prep Kit with Ribo-Zero Gold Kit (Illumina). These libraries were sequenced using Illumina HiSeq platform with 100 bp single read at a depth of 10-20 million reads per sample.

### Bioinformatic analyses of bulk RNA-Seq

RNA-Seq reads were demultiplexed using bcl2fastq2, aligned to UCSC genome mm10 with STAR aligner ^50^ and Ensembl 94 gene models. TPM values were generated using RSEM. Downstream analysis was performed in R using the read counts generated by STAR aligner and the TPM values. Differential expression analysis was performed using DEseq2^51^ with likelihood ratio test. The TPM values of the differentially expressed genes were used for visualization. Differentially expressed genes were clustered into 3 groups using Ward.D2 method based on gene wise Pearson correlation. Metagene was plotted as the locally estimated scatterplot smoothing (LOESS) regression of the genes in the same group at the same time point. GO term analysis was performed using “goseq” package.

### scRNA-Seq of the olfactory epithelium

The P0, P3, P7, and P21 data were from the published dataset ^23^. Data of three new time points, E18.5, P14, and adult, were acquired with the same method and added to the analysis. Briefly, the OE from CD-1 mice were dissected in oxygenated artificial spinal cord fluid (ACSF). Single cells were dissociated with papain at 37□ and filtered using 10 µm pluriStrainer (PluriSelect).

Filtered cells were stained with DAPI and Draq5. DAPI negative, Draq5 positive nucleated live single cells were sorted using BD Influx cell sorter. scRNA-Seq was performed using 10X Chromium single cell platform (10X Genomics). Libraries were prepared using Chromium Single Cell 3’ Library & Gel Bead Kit v2. Libraries were sequenced using Hi-seq platform with ∼50 thousand reads per cell.

### Bioinformatic analysis of scRNA-Seq data

The scRNA-Seq data were processed using Cell Ranger pipeline to acquire a filtered UMI count matrix. This matrix was further analyzed in Seurat ^52^ with the following steps. Genes expressed by more than 5 cells were used. The cells were first filtered with genes per cell between 1500 and 5000. The percentage of mitochondria genes was set to be lower than 2.5%. The data was then normalized through scTransform procedure using “SCTransform” function. Principal component analysis was performed to extract latent variables and reduce the dimensions. Top 20 principal components were used for unsupervised clustering. Cell types were identified by constructing a Shared Nearest Neighbor (SNN) Graph using “FindNeighbors” function. The clusters were then determined by optimizing the modularity function using “FindClusters” function. Specifically, the Clusters were assigned to cell types by the expression of canonical cell markers. Globose basal cells, intermediate neuronal progenitor cells, iOSNs, and mOSNs were identified by the expression of Ascl1, Neurog1, Lhx2, Hdac2, Gap43, and Omp for downstream analysis. The cells were plotted in 2 dimensions by UMAP algorithm using “RunUMAP” function. To compare the gene expression pattern betweenRNA-Seq, and scRNA-Seq, scTransform normalized gene counts from the OSNs of the same age samples were averaged and hierarchically clustered into three clusters.

### SLIDE-Seq

The SLIDE-Seq2 puck used in this study was a gift from Dr. Fei Chen prepared according to the methods described previously^29^. The puck was stored in the dark at 4□ prior to use. OE tissue dissected from P3 CD1 mouse was embedded in optimal cutting temperature compound (O.C.T., Sakura Finetek), snap-frozen in liquid nitrogen, and stored at -70°C until use. OE tissue was sectioned into 10 µm and mounted onto the puck using a cryostat (CryoStar NX70). The puck was placed into a 1.5 mL tube for library preparation. The puck was immersed in 200 µL of hybridization buffer (6X SSC, 2 unit / µL Lucigen NxGen RNAse inhibitor) for 30 minutes at 37°C for the binding of RNA to the oligos. First strand synthesis was performed in RT solution (75 µL water, 40 µL 5X Maxima RT buffer, 40 µL 20% Ficoll PM-400, 20 µL 10 mM dNTPs, 5 µL RNase Inhibitor, 10 µL 50 M Template Switch Oligo, 10 µL Maxima H^-^ RTase) for 1 hour at 42□. After reverse transcription, tissue was removed by adding 200 µL 2X tissue digestion buffer (200 mM Tris-Cl, pH 8, 400 mM NaCl, 4% (w/v) SDS, 10 mM EDTA, 32 unit / µL Proteinase K) and incubation at 37□ for 30 minutes. Beads were removed from the slide by pipetting. Beads were then washed in 200 µL wash buffer (10mM Tris pH 8, 1 mM EDTA, 0.01% Tween-20) for 3 times, washed with 200 µL water, and resuspended in PCR mix (22 µL water, 25 µL Terra PCR direct buffer, 1 µL Terra Polymerase (Takara), 1 µL 100 M Truseq PCR handle primer, 1 µL 100 µM SMART PCR primer). Library was amplified by PCR. PCR product was purified with Ampure XP beads (Beckman Coulter), resuspend in 20 µL water, and quantified with Bioanalyzer (Agilent). 600 pg of PCR product was used to generate Illumina sequencing library using Nextera XT kit (Illumina). The library was sequenced on a high output flow cell using Illumina NextSeq 500. A 75 cycle High Output kit v2 was used with the following paired read lengths: 42 bp Read 1, 8 bp i7 index, and 41 bp Read 2.

### Doxycycline treatment

To inhibit transgene expression from tetO promoter, nursing CD1 dams were fed with DOX containing chow (Envigo) at least 48 hours before the experiment. Pups were fostered with the dams to stop transgenic expression. DOX diet was maintained until the animals were sacrificed unless otherwise stated. To induce transgenic Fzd1 expression from tetO promoter, breeding *Fzd1*^*EE*^ animals were fed with DOX containing chow when paired and the pups were fostered with CD1 dams fed with regular chow at the indicated time points to remove repression by DOX.

