## supplemental material for "Molecular Control of Circuit Plasticity and the Permanence of Imprinted Odor Memory"

### Extended Data

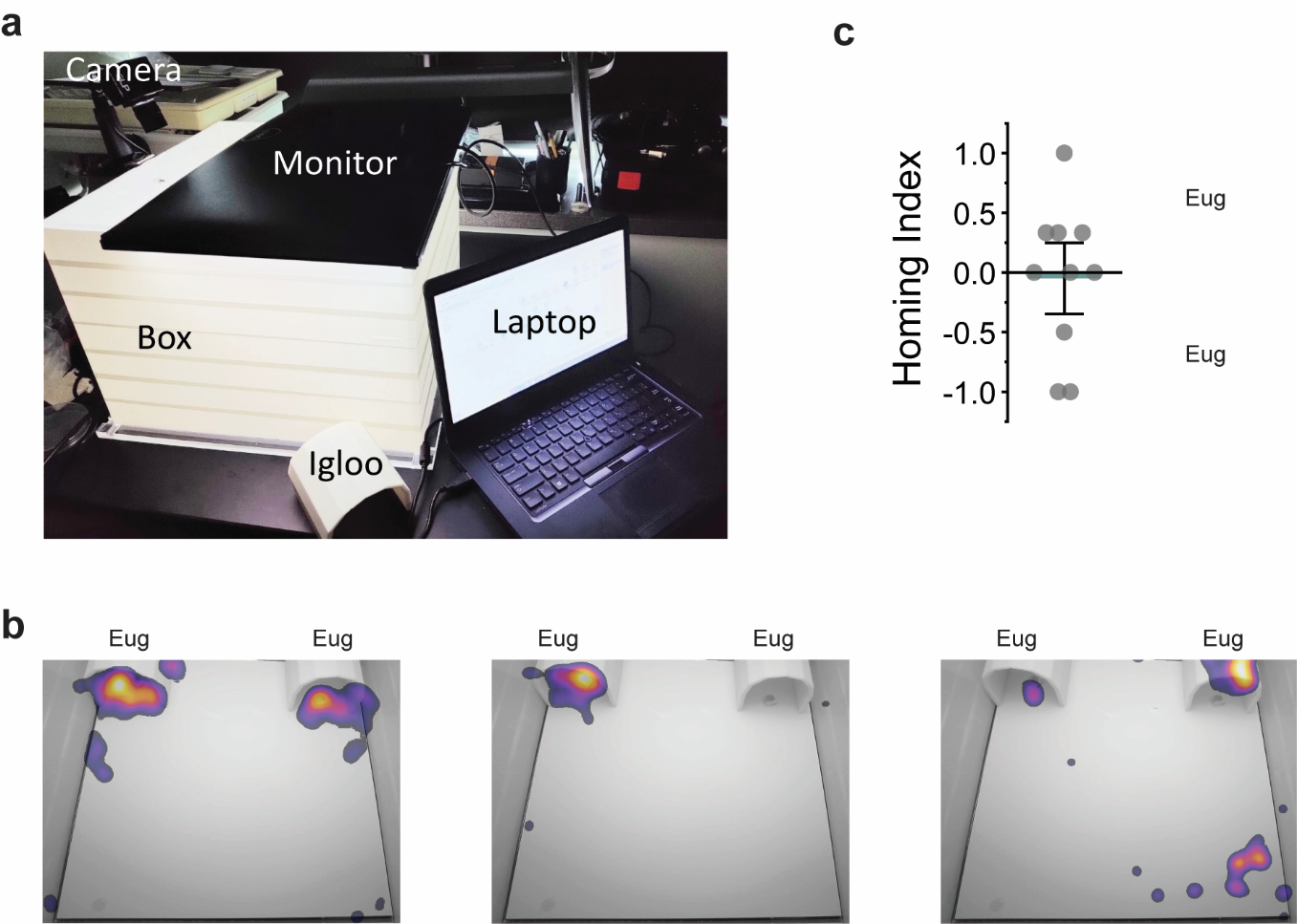

**Fig. 1. Threat induced homing responses assay.** **a**. Photograph of the TIHR set up. **b**. Cumulative heatmaps of dwell time following looming threat for three animals when eugenol odor is placed in both igloos. Animal on the left hide under the two igloos with equal probability. The other two animals consistently opt for one side to hide. **c**. Quantitative data when both igloos have eugenol odor.

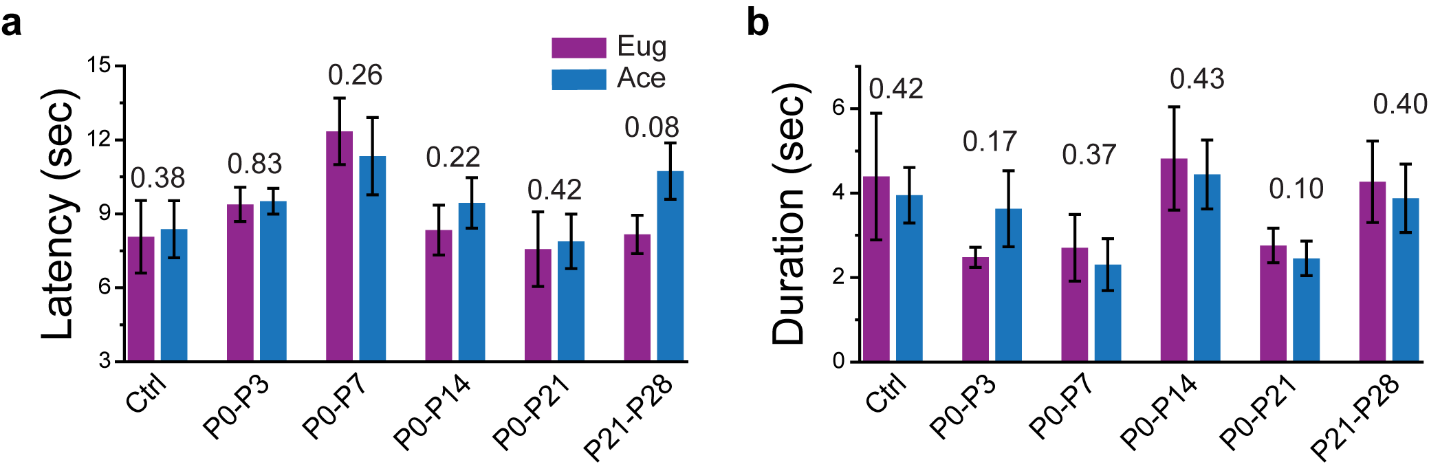

**Fig. 2. Detailed analysis of TIHR assay.** **a**. Latency to escape for the animals tested. **b**. Duration of escape for the animals tested. P-values are indicated above the bars.

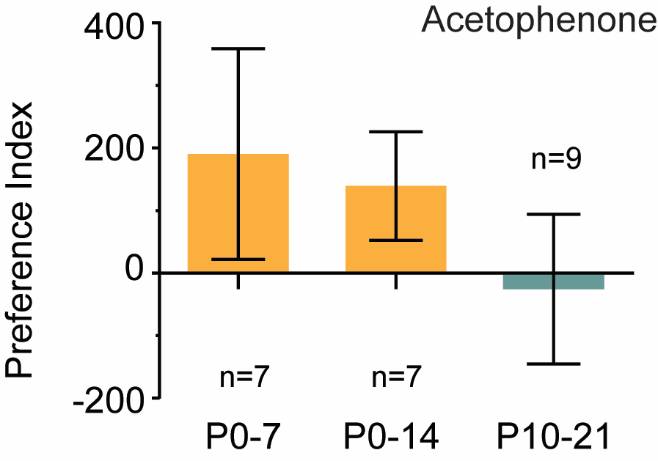

**Fig. 3. Mice maintain the same odor preferences after six months.** Mice tested in Fig. 1 were tested at 6 months.

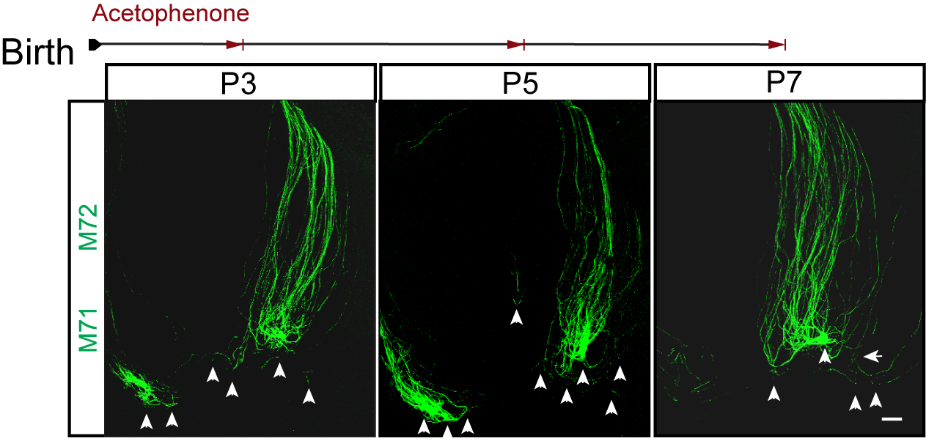

**Fig. 4. Odor induced changes in axon projection of OSNs expressing cognate receptors.** Neonatal M71G;M72G pups were exposed to acetophenone, and their dorsal OB imaged at P3, P5, and P7. Arrowheads indicate individual glomeruli containing GFP labeled axons determined from 3-D images. Scale bar: 100 µm.

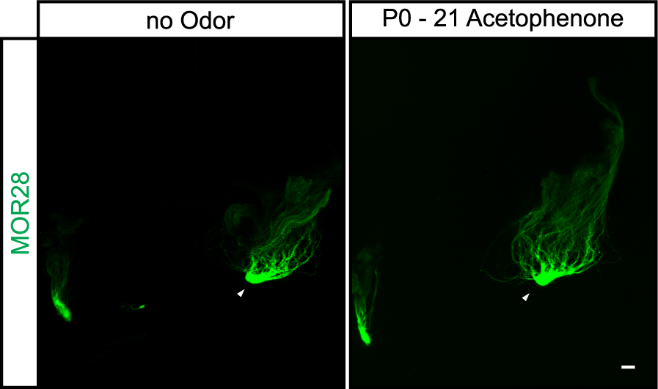

**Fig. 5. Odor does not induce changes in axon projection of OSNs expressing other receptors.** Neonatal MOR28-GFP pups were exposed to acetophenone, and their dorsal OB imaged at P21. Arrowheads indicate individual glomeruli containing GFP labeled axons determined from 3-D images. Scale bar: 100 µm.

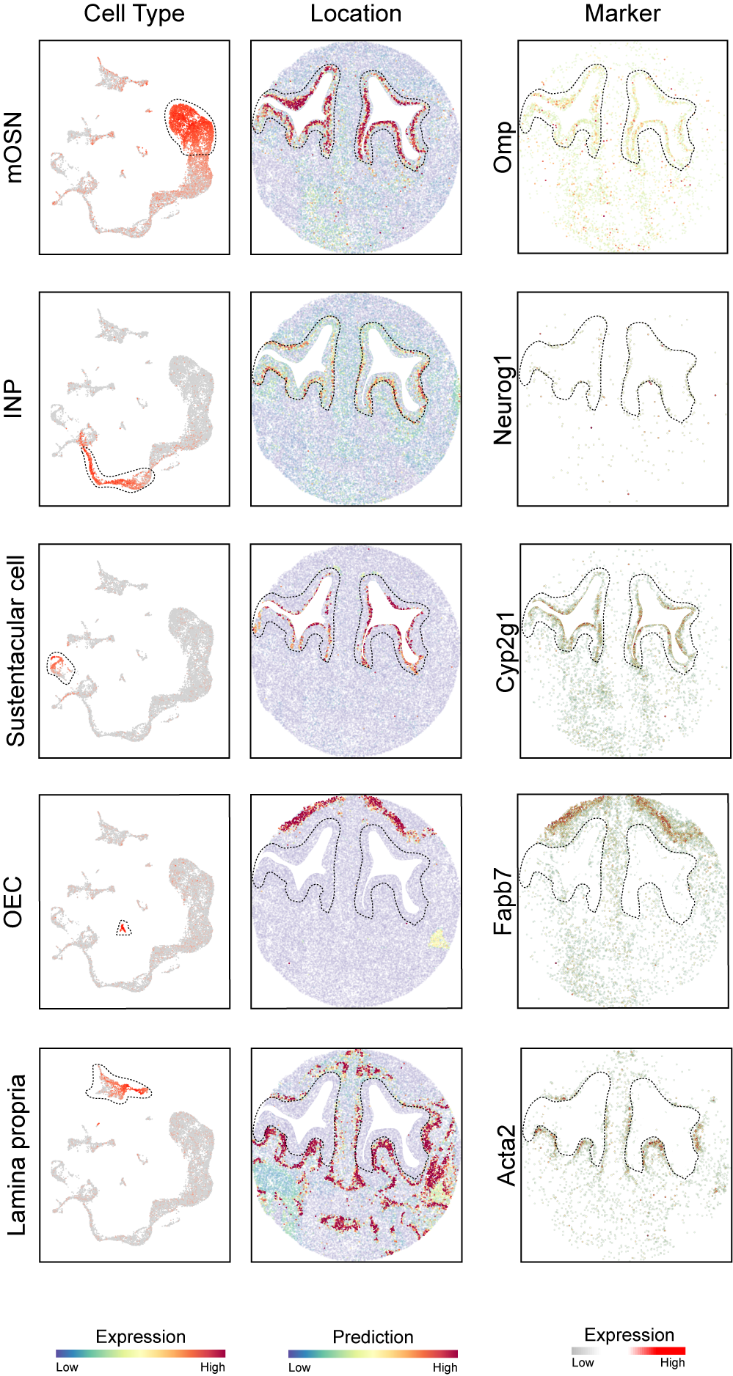

**Fig. 6. Slide-Seq localization of different cell types.** Left panel, cell types identified by scRNA-Seq. Middle panels, spatial location of cell types in Slide-Seq. Right panels, expression patterns of marker genes in scRNA-Seq. Gene expression level presented as SCT.

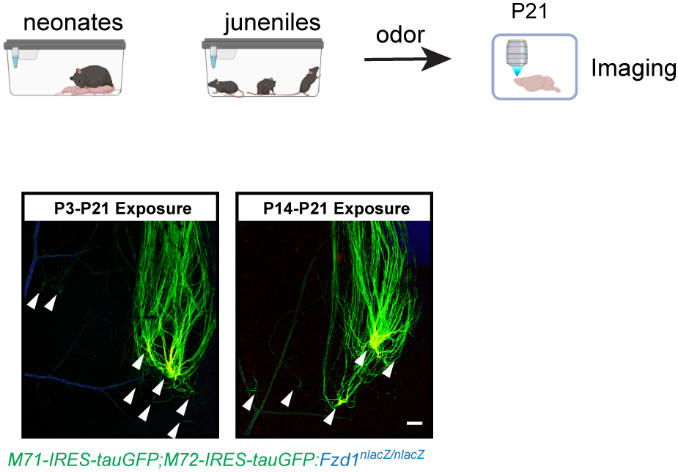

**Fig. 7. Odor induced changes in axon projection in Fzd1 KO mice.** Top: schematic of experimental design. Bottom**:** M71G;M72G; *Fzd1^nlacZ/nlacZ^* pups were exposed to acetophenone starting at P3 (left) or P14 (right) and their dorsal OB imaged at P21. Compared to Fig 2 b and c, odor exposure starting at P14 still induced broadened projection patterns. Arrowheads indicate individual glomeruli containing GFP labeled axons determined from 3-D images. Scale bar: 100 µm.

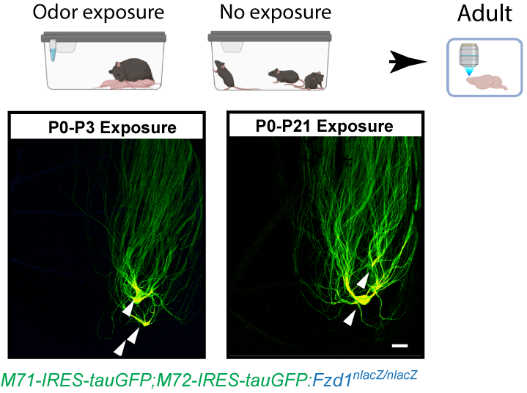

**Fig. 8. Recovery of axon projection following odor removal in Fzd1 KO mice.** Top: schematic of experimental design. Bottom**:** M71G;M72G; *Fzd1^nlacZ/nlacZ^* pups were exposed to acetophenone starting at P0 with the odor removed at P3 (left) or P21 (right) and their dorsal OB imaged at the adult stage. Compared to Fig 2 e and f, axons continued to form convergent projection even when odors were removed at P21 Arrowheads indicate individual glomeruli containing GFP labeled axons determined from 3-D images. Scale bar: 100 µm.

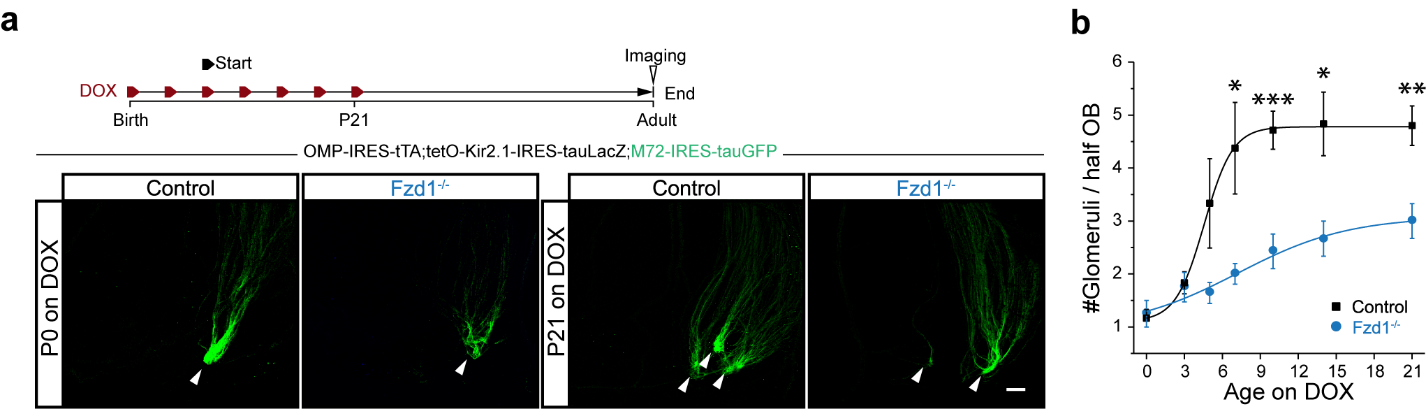

**Fig. 9 Extended plasticity in** *Fzd1^nlacZ/nlacZ^* **mice. a.** Projection patterns of M72 axons in control and *Fzd1^nlacZ/nlacZ^* mice that carry the *M72-IRES-GFP;Omp-IRES-tTA;tetO-Kir2.1* compound alleles. Upper panel, schematic illustrating DOX given at different time points between P0 to P21 to stop Kir2.1 expression and restore neural activity. Bottom panels, representative images of M72 glomeruli (green) from *Fzd1^nlacZ/nlacZ^* and their littermate control animals. Axon projections were imaged at the adult stage. Arrowheads indicate individual glomeruli. **b**. Quantification of the experiments. Control, n = 10, 12, 6, 8, 7, 6, and 5 for P0, P3, P5, P7, P10, P14, and P21. *Fzd1^nlacZ/nlacZ^*, n = 8, 12, 14, 15, 7, 6, and 16. P = 0.5907, 0.8127, 0.1027, 0.0289, 0.0001, 0.0140, and 0.0065.

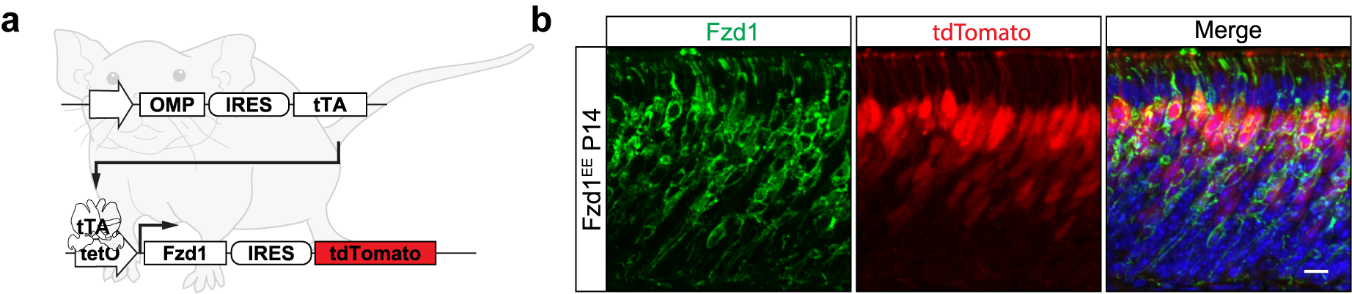

**Fig. 10 Ectopic expression of Fzd1. a.** Illustration of using tetOff strategy to express Fzd1 in the OSNs in OMP-IRES-tTA;tetO-Fzd1-IRES-tdTomata (*Fzd1^EE^*) mice. **b**. Detection of Fzd1 protein (green) and tdTomato (red) in the OE of the *Fzd1^EE^* mice.

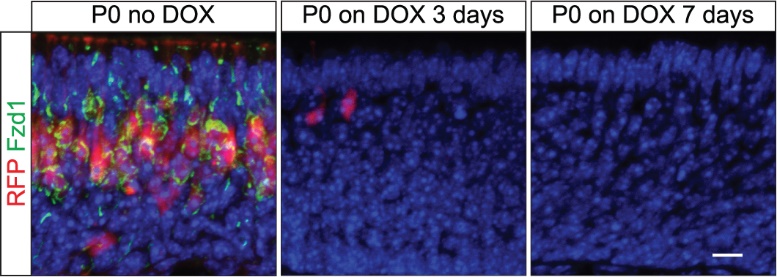

**Fig. 11 DOX effectively turning off ectopic expression of Fzd1.** Fzd1 (green) and tdTomato (red) expression in the olfactory epithelia of the *Fzd1^EE^* mice with no DOX (left), DOX from P0-P3 (middle), or from P0-P7.

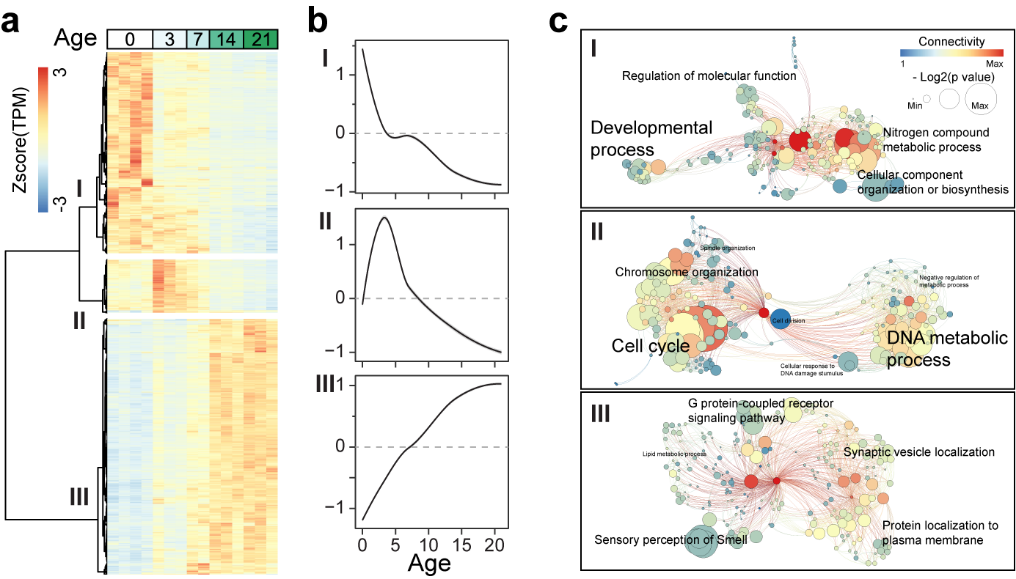

**Fig. 12 Transcriptome change of the OE during postnatal development. a.** Heatmap of differentially expressed genes in the OE between P0 and P21 detected in RNA-Seq (n = 4, 3, 2, 3, and 3 for P0, P3, P7, P14, and P21). Data is presented as Z-score of log2 (TPM + 1). **b**. Metagene expression analysis for the three types of genes from (a). Metagene is calculated as a locally estimated scatterplot smoothing (LOESS) of genes within the group. **c**. GO term enrichment analysis for the three groups of genes. Size of the circle indicates -log2 of the p value of the GO term. GO terms are connected according to their hierarchy. Color indicates the number of parents and offspring the term has.

**Movie 1**. TIHR assay for a mouse without exposure to acetophenone. The two igloos have pads with eugenol (left) and acetophenone (right) odors, respectively.

**Movie 2**. TIHR assay for a mouse which exposed to acetophenone at P0-14. The two igloos have pads with eugenol (right) and acetophenone (left) odors, respectively.
